# Neurophysiological and Behavioral Responses of *Ixodes scapularis* to host odors

**DOI:** 10.1101/2020.08.04.236539

**Authors:** Tanya Josek, Jared Sperrazza, Marianne Alleyne, Zainulabeuddin Syed

**Affiliations:** Center of Mathematics, Science, and Technology, Illinois State University, Normal, IL 61790, USA; Department of Entomology, University of Kentucky, Lexington, KY 40546, USA; Department of Entomology, University of Illinois at Urbana-Champaign, Urbana, IL 61801, USA

**Author notes:** Corresponding author: Zainulabeuddin Syed, Department of Entomology, University of Kentucky, Lexington, KY 40546. **Email:**.

**Keywords:** Single sensillum recordings, GC-linked single sensillum recordings, Dual-choice assay, GC-Mass spectrometry, Blacklegged tick

## Abstract

The black-legged tick, *Ixodes scapularis* (Ixodida, Ixodidae), is one of the major disease vectors in the United States and due to multiple human impact factors, such as decreasing forest size for land development and climate change, it has expanded its range and established across the United States. Throughout the life cycle, ticks locate hosts for their blood-meal and although the ecologies of this tick and their hosts have been studied in depth, the sensory physiology behind host location largely remains unexplored. Here we report establishing a robust paradigm to isolate and identify odors from the natural milieu for *I. scapularis*. We performed single sensillum recordings (SSR) from the olfactory sensilla on the tick tarsi, and used the SSR system as biological detector to isolate natural compounds that elicited biological activity. The SSR setup was further tested in tandem with gas chromatography (GC) wherein the ticks’ olfactory sensillum activity served as a biological detector. The GC-SSR recordings from the wall pore sensilla in the Haller’s organ, and further identification of the biologically active deer glad constituents by GC-mass spectrometry (GC-MS) revealed methyl substituted phenols as strong chemostimuli, as compared to ethyl or propyl substitutions. Strongest electrophysiological activity was elicited by *meta-cresol* followed by *para*-cresol. Ethyl- and propylphenols with any of the three, *ortho, meta* or *para* substitutions, did not induce any neurophysiological activity. Finally, a behavioral analysis in a dual-choice olfactometer of all these phenols at three different doses revealed no significant behavioral response, except for *p*-cresol at −3 dilution Overall, this study contributes to our understanding of *I. scapularis* tick’s neurophysiology and provides a robust platform to isolate and identify natural attractants and repellents.

## INTRODUCTION

Ticks, like all other blood feeding arthropods, display robust olfactory driven behaviors (Allan 2010, Mulenga 2013, Syed 2015). However, unlike insects where antennae serve as primary olfactory organs, ticks use their first pair of legs for odor detection. A distinct apparatus called the Haller’s organ is located on the foretarsal region of the first pair of legs and was originally assumed as an hearing organ (Haller 1881). A direct role of this organ in olfaction has since been established in multiple members of tick species (Carr et al. 2017). Additional functions ascribed to the Haller’s organ are the detection of *liquid* water (Krober and Guerin 1999), and infrared detection (Mitchell et al. 2017, Carr and Salgado 2019). Detailed ultrastructural analyses in hard- (Hess and Vlimant 1986) and soft-ticks (Klompen and Oliver Jr 1993), and more recently from the field collected ticks (Josek et al. 2018a) have shown multiple olfactory sensilla that are present in and around the Haller’s organ. Host location and attachment in *Ixodes scapularis* begins with their climbing up a grass blade or shrubs and waiting for the approaching host animal as they raise up their forelegs (Nicholson et al. 2019). *I. scapularis* feed on multiple animals species, even within one life cycle, such as white-tailed deer, white-footed mice, chipmunks, or raccoons (LoGiudice et al. 2003). Because *I. scapularis* prefers different hosts during different life stages (Bishopp and Trembley 1945, Schulze et al. 1986, LoGiudice et al. 2003, Keesing et al. 2009), it is conceivable that they detect a variety of odors (Allan 2010).

In blood-feeding arthropods a distinct yet limited range of volatiles from the environment are parsimoniously used in various contexts eliciting distinct behaviors such as *attraction* to host and mates, and *repulsion/avoidance* from unsuitable sites (Syed 2015). These behavior modifying chemicals (semiochemicals) are detected by the olfactory receptor neurons (ORNs) that innervate the sensilla in the Haller’s organ (Hess and Vlimant 1986). Determining the detection properties of these ORNs is critical in isolating and identifying the volatile chemicals that will lead to the development of odor baits to be used in tick management programs. The single sensillum recording method (SSR) is one of the most robust techniques used to record from individual ORNs. When using SSR each ORN is challenged with a variety of semiochemicals and the induced responses are recorded either as excitation or inhibition. The method has widely been used in insect studies (Kaissling 1995, Olsson and Hansson 2013). The SSR can also be coupled with gas-chromatography (GC-SSR) to isolate and identify biologically active constituents from the complex odors that stimulate ORNs in blood-feeding arthropods (Reisenman et al. 2016). The GC-SSR protocol has only been used in limited studies in ticks due to the difficulty in securing stable recordings from Haller’s organs that are located in strong forelegs. The first and only GC-SSR recordings from *Amblyomma variegatum* identified a range of chemostimuli from host odors (Steullet and Guerin 1992b, a, 1994a, b), a preliminary study in *I. ricinus* identified mostly phenolic compounds as eliciting a response (Leonovich 2004), and in *Boophilus microplus* 2,6-dichlorophenol was isolated and identified as the key electrophysiologically active constituent from the extracts of male, female and larva (De Bruyne and Guerin 1994). Here we report the first GC-SSR recordings from *I. scapularis* that identified a range of electrophysiologically active constituents that were further evaluated for the behavioral responses from adult.

The blacklegged ticks have doubled their distribution range in the past 2 decades (Eisen and Eisen 2018). Also, at least 8 infective microorganisms vectored by ticks have established during this period thus warranting tools for reliable and robust monitoring and control strategies. Multi-pronged management strategies that explore and expand on innovative control methods, such as exploitation of vectors’ strong sense of smell provide effective practices that can be embedded into the existing vector control strategies (Allan 2010, Carr and Roe 2016, Esteve-Gassent et al. 2016). Odors, such as pheromones, are a proven means to sample and control vector arthropods (Pickett et al. 2010), including ticks (Sonenshine 2006), yet identifying an effective and selective bait has been elusive so far (Syed 2015, Carr et al. 2017). Knowing how these cues are detected and perceived by ticks will help us understand their role in the tick’s life cycle and aid in maximizing the effectiveness of management strategies.

## METHODS

### Ticks and Tick Maintenance

*Ixodes scapularis* ticks were obtained from the Oklahoma State University tick rearing facility and from the Centers for Disease Control and Prevention for distribution by BEI Resources, NIAID, NIH (NR-42510). Ticks were maintained at 22 ± 2°C, 99% RH and 15:9 h light/dark cycle. Only adult female ticks were used in experiments.

### Electrophysiology

#### Stimuli

A set of natural and synthetic stimuli were tested: Commercially available preorbital gland extracts (Jackies Deer Lures^®^), deer tarsal gland extracts (Harmon Scents^®^), deer interdigital gland extracts (Harmon Scents^®^) were obtained. Synthetic CO_2_ was obtained from Airgas, USA. In addition to natural extracts and CO_2_, mixtures of synthetic chemicals (phenol, carboxylic acids, aldehyde, or alcohol; all potential host semiochemicals) suspended in hexane were tested at 10^-2^ dilutions (Table 1). Following the consistently excitatory responses elicited by the phenol mixture, we prepared multiple sub-mixtures of phenols: each of the three mixtures included phenol supplemented with either of the three *alkyl* phenols of increasing chain lengths (*methyl*-, *ethyl*- and *propyl*-phenol). Therefore, phenol mixture 1, 2 and 3 denote each of this mixture composed of phenol added to the either of the three *alkyl* phenols in *ortho, meta* or *para* configurations respectively. The stimuli, unless otherwise mentioned, were purchased from Sigma-Aldrich (St Louis, MO) and were all above 98% pure.

**Table 1:** Contents of the odor mixes used for SSR and induces responses. o: No response; + is up to 20% increase; ++ is 25–50% increase, and +++ is 50–100% increase. Percentage increase is determined after the maximum percentage increase of the excitation response from the ORNs in the sensillum upon stimulation with - 2 dilution of *m*-methylphenol (*ca*. 250 spikes/s)

| Name of the stimuli | Contents | Response |
| --- | --- | --- |
| CO<sub>2</sub> | Gaseous carbondioxide ~5% (Airgas) | o |
| Tarsal gland | Commercial Extract (Harmon Scents®) | ++ |
| Interdigital gland | Commercial Extract (Harmon Scents®) | ++ |
| Preorbital gland | Commercial Extract (Jackies Deer Lures®) | ++ |
| *o* - Phenol Mixture | 2-methylphenol, 2-ethylphenol and 2-propylphenol | + |
| *m* – Phenol mixture | 3-methylphenol, 3-ethylphenol and 3-propylphenol | +++ |
| *p* – Phenol mixture | 4-methylphenol, 4-ethylphenol and 3-propylphenol | ++ |
| Carboxylic Acid mixture | propionic acid, 2-methylpropanoic acid, butyric acid, 2-methylbutanoic acid, acid, pentanoic acid, hexanoic acid and DL- lactic acid | ++ |
| Alcohol mixture | butanol, pentanol, hexanol, heptanol, octanol, nonanol, 3-methylbutanol, 2-butoxyethanol, octanol, 2-hexen-1-ol, *E*-3-hexen-1-ol, and *E*-2-hexen-1-ol | o |
| Aldehyde mixture | propanal, pentanal, heptanal, nonanal, butanal, hexanal, octanal, and decanal | + |

#### Stimulation

Initially the natural extracts, CO_2_ and synthetic mixtures were screened at least three times on a given tick preparation. A typical offline SSR recording (not linked to GC) lasted for 10 s consisting of a 1 s pre-stimulus period, a 500 ms pulse of the odor, and 8.5 s post-stimulus recording. Odors that induced excitatory responses from the sensillum were further tested by GC-SSR. This whole procedure was repeated with at least three females for each odor tested.

GC-linked SSR method have been described elsewhere (Steullet and Guerin 1992b, Syed 2015). In short, ticks were immobilized on ice. Adult females were mounted on a microscope slide using double sided adhesive tape (3M Scotch, USA) ventral side down. The body was placed on the tape first and then, using forceps, legs were gently spread on the tape. The forelegs were stretched forward and orientation was maintained in such a way that the Haller’s organ leg was facing up. The tick was subsequently completely covered, except for the front two pairs of legs, in G-U-M^®^ unscented dental wax (Sunstar Americas, Inc., USA) (Figure 1A). The tip of the second leg on the right side of the tick was excised with surgical microscissors (World Precision Instruments, Sarasota, FL) and connected with an indifferent electrode. Electrodes contained silver wires in drawn-out glass capillaries filled with 0.1% KCl and 0.5% polyvinylpyrrolidone (PVP) (Sigma-Aldrich, USA). The tick preparation with the ground electrode impaled and held in position was moved under a high magnification microscope (Olympus BX51WI) that was continuously bathing in a clean and humidified airflow (Scheidler et al. 2015). A recording electrode was maneuvered with a MPM-10 Piezo Translator (World Precision Instruments, USA) to the base of the sensillum to make a firm contact with the wall pore (wp) sensillum inside the capsule aperture of the Haller’s organ (Josek et al. 2018a) (Figure 1B).

**Figure 1:**
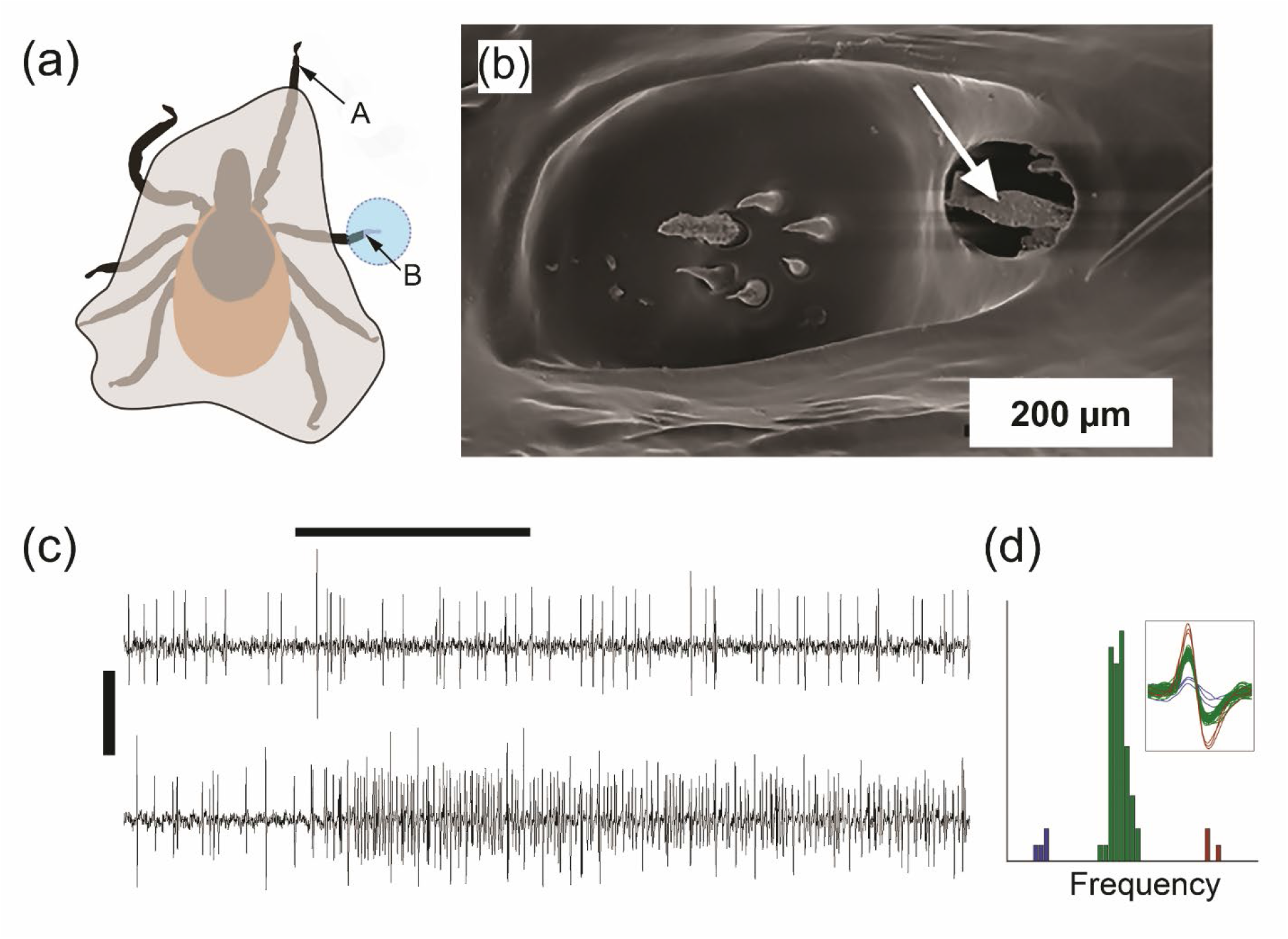
Single unit electrophysiological recordings set-up for recording from *Ixodes scapularis* (a) Sketch of the restrained tick. Dental wax, highlighted in grey with a solid surrounding line, completely covers all but the front two pairs of legs. A-Recording region i.e., foreleg with the Haller’s organ exposed, and B is the location of the ground electrode placement. (b) Environmental scanning electron microscope image of the Haller’s organ of *I. scapularis*. White arrow indicates the wall pore sensillum used for the recordings, (c) Extracellular single-unit recordings from a the sensillum reveal the presence of 3 ORNs with distinct response characteristics. *Upper trace* is spontaneous activity showing 3 neurons of specific amplitude and frequency. *Lower trace* indicates excitatory responses to *o-c*resol (−3 dilution). Scale bars are 500 (horizontal) and 0.5 mV (vertical), and (d) amplitude-frequency histograms showing 3 distinct neuronal spikes clusters.

Signals were amplified and directly recorded using a analogue-digital IDAC4-USB box (Syntech, Germany) via a high-impedance preamplifier and recorded on the hard disk of a PC. Recordings were analyzed with the AutoSpike V3.9 software (Syntech). AC signals (APs or spikes) were band-pass filtered between 100 and 10,000 Hz, and for the DC signals (receptor potentials/sensillum potentials,SPs), a high filter of 3 kHz was used. The activity of co-located ORNs in a sensillum could be assessed based on differences in spike amplitude and frequency (Figure 1 C & D). However, in this work, we report the activity in terms of percent activity change that represents sum of the spikes generated from all the ORNs in the sensillum upon stimulation. Resolved odor constituents from the GC column were added into the main flow. GC-SSR was performed on a GC-7890 Agilent Gas Chromatogram equipped with a DB-5 column (30 m, 0.25 mm; 0.25 μm; Agilent Technologies) and connected with a transfer line and a temperature control unit from Syntech, Germany. Effluents from the capillary column were split at the three way splitter connected with make-up gas (Agilent) into two fractions with 2:1 ratio. The major fraction was directed to the continuous airflow bathing the sensilla, and the other fraction to the flame ionization detector. Later, extracts and synthetic mixtures were injected onto the comparable column in an Agilent 5975C Series GC-MS with Triple-Axis Detector. Biologically active constituents were identified by 3 criteria: 1-matching spectra with those in the NIST 2008 MS library; 2-Kovat’s Retention Indices (KI), and 3-confirming the biological activity by GC-SSR. The program for GC-SSR and GC-MS was the same: starting at 5Ü°C holding for 1 minute then ramping 10°C/minute to 3°C followed by a hold of 5 minutes. Flow was 3 ml/min for GC-SSR and 1 ml/min for GC-MS.

In order to determine the change in the signal after exposure to the odor, we first determined the delay in odor exposure or the time it takes an odorant to travel from GC to the tick preparation. The delay time was used to set a time period to count action potentials prior to and during the stimulation. The percent changes in spike rates were then calculated. Analyses and graphical representations of these data were created using the packages “stats”,“dplyr”, “tidyr”, “Rmisc”, and “ggplot” in R version 3.5.1 (R core team, 2017) within Rstudio environment version 1.1.463 (Rstudio Team, 2016). Data collected were first checked for normality with the Shapiro-Wilk Normality Test (Shapiro and Wilk 1965). A pairwise t-test with a Holm *p*-adjustment method (Holm 1979) was used to determine if the percent changes in spikes were significant.

### Behavioral Choice Tests

The behavioral choice assay was created based on the original model (Geier and Boeckh 1999) with a few modifications. This dual-choice olfactometer was designed and fabricated using laser cut acrylic (Figure 4A&B). Air entered the olfactometer through two separate points each at a flow rate of 40 ml/min and was pulled out using a vacuum at 80 ml/min. Air was pushed into and out of the olfactometer using a the portable PVAS22 Volatile Assay System (Rensselaer, New York, USA). Each air flow tube had a cloth mesh preventing ticks from entering the tubes. Prior to entering the olfactometer, the air was humidified by drawing the air through a 150 ml Erlenmeyer flask partially filled with dH_2_O; on the experimental side air also passed through a vial containing a 4 x 1 cm piece of filter paper treated with test solutions. We tested all the alkyl constituents from the three phenol mixtures. A 0.5 ml of the phenol mixture (at 10, 100 or 1000x dilution in ethanol, represented here as −1, −2 and −3 dilutions, respectively) were used as test stimuli. Ethanol (0.5 ml) applied to a filter paper served as control. For each trial, four starved adult ticks were randomly selected and allowed 1 min in the olfactometer stem to acclimatize. Ticks were given a maximum of 3 minutes to make a choice. Odor induced responses were recorded as a choice arm if a tick crossed a choice after passing a designated line (~7cm from the ticks release point) placed on each branch of the olfactometer (Figure 3A). This was repeated until a total of twenty trials, totaling 40 males and 40 females, were conducted. Ticks were only used once per treatment type to avoid pseudoreplication. Treatments were randomized and the olfactometer was rinsed with 100% ethanol and allowed to dry for 30 min between trials.

Data were analyzed using the ‘stats’ and ‘FSA’ packages in R version 3.5.1 (R core team, 2017) within Rstudio environment version 1.1.463 (Rstudio Team, 2016) in order to determine if there were differences in overall activity, if any choice was made (towards either side) versus no choice being made. Data were checked for normality using the Shapiro-Wilk Normality Test (Shapiro and Wilk 1965). A Kruskal–Wallis Rank Sum Test (Kruskal and Wallis 1952) was used to check differences within the means of samples. Pairwise comparisons were made using a Dunn test (Dunn 1961) and Holm *p*-adjustment method (Holm 1979). A Wilcoxon signed-rank test (Wilcoxon 1945) with Holm *p*-adjustment method (Holm 1979) was used in order to compare tick choice for each treatment type.

## RESULTS

Single unit recordings from the Haller’s organ sensillum by our preparation method (Figure 1A) were stable and highly reproducible. The extracellular action potentials from the ORNs inside the large sensillum in the capsule aperture (Figure 1B) revealed at least 3 ORNs that were spontaneously active and were clearly distinguishable by their amplitude and frequency (Figure 1C-D). Commercial odors derived from the most preferred host of black-legged tick, white tail deer, and a broad array of arthropod semiochemicals (Syed 2015) induced a variety of responses. Synthetic mixtures of phenols, aldehydes and carboxylicacids induced excitatory responses, whereas alcohols inhibited the spontaneous activities of all the three ORNs for a short period of time during the odor exposure (Table 1). Following the observations that the commercially available deer gland extracts induced high excitatory responses, we analyzed them on GC-MS to identify their chemical composition. All the three extracts were rich in phenols: phenol, *meta*- or *para*-cresol, and *meta*- or *para*-ethylphenol were major constituents by abundance (Supplemental Figure 1). We determined the Kovat’s Retention Indices (KI) for these compounds as 988, 1080 and 1168 respectively (from left to right) on DB-5 column. Since the DB-5 column could not resolve between the *meta* or *para* configuration, we systematically prepared synthetic phenolic mixtures of all the three configurations of varying chain lengths and analyzed them by GS-SSR.

Of all the alkyl phenols evaluated by GC-SSR, consistently stronger responses were induced by methylphenols (Figure 2A and 3). Significantly higher responses were recorded to *meta*- and *para*-configurations of methylphenols (Figure 3). Methyl, ethyl or propylphenols in any other configurations did not elicit any significant increases in overall spikes from the sensillum over the spontaneous activity.

**Figure 2.**
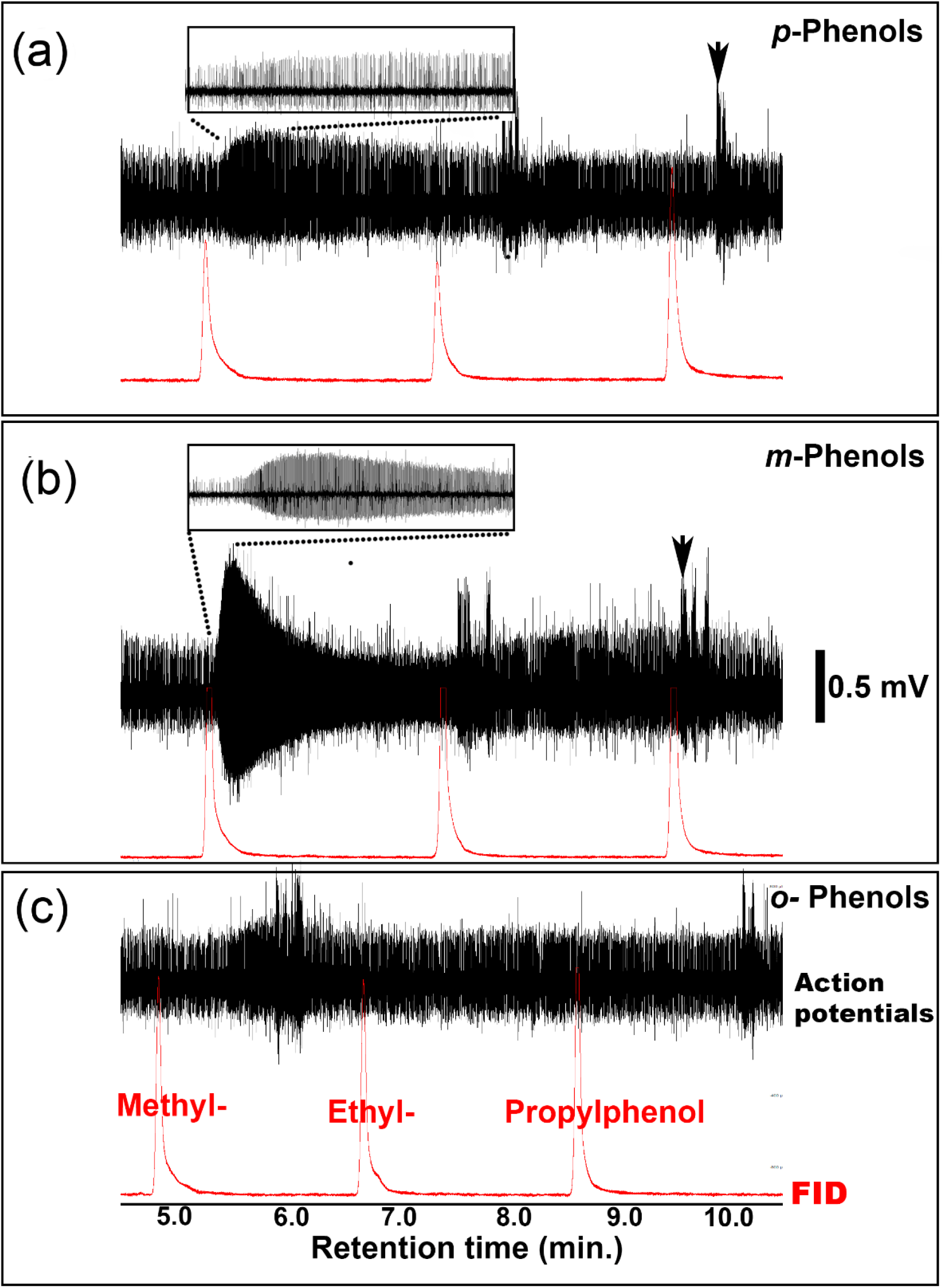
Analysis of phenolic mixtures by gas chromatography-linked electrophysiology (GC-SSR) from the sensillum within Haller’s organ in a female *I. scapularis*. The mixtures analyzed are composed of (a) methyl- (b) ethyl- or (c) propylphenols. Note the comparable elution time in (a) and (b); *ortho* phenols (c) elute at slightly earlier time. Expanded traces (dotted rectangles) in (a) and (b) depict the excitation patters; maximum response was elicited by the *m*-methylphenol followed by *p*-methylphenol. Occasional bursts indicated by the arrows heads represent strong muscle potential due to occasional leg movements. The bottom red trace in each of the 3 recordings is the flame ionization detector (FID) response, and upper traces are real time action potentials.

**Figure 3.**
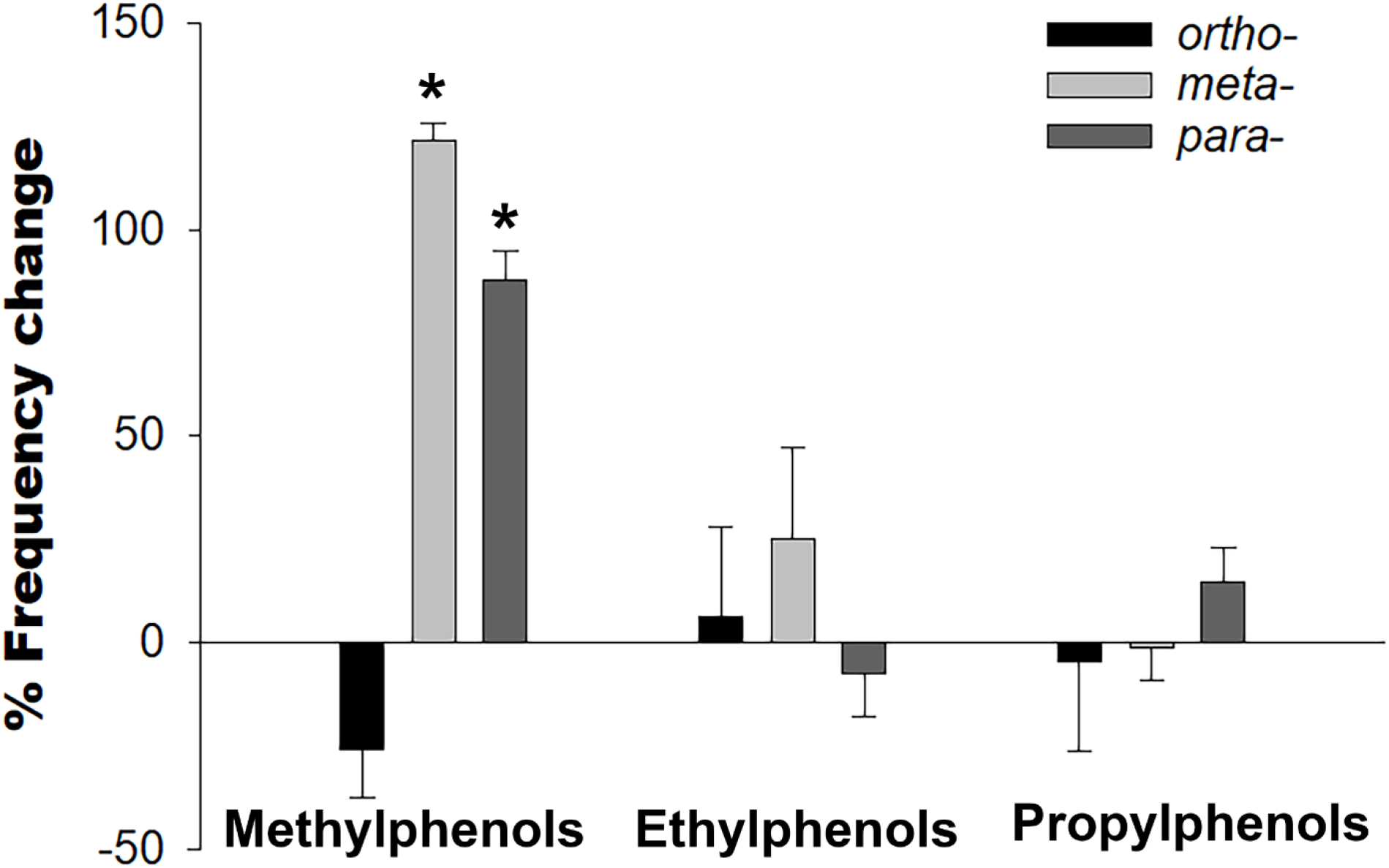
Electrophysiological response elicited by alkyl phenols (data from Figure 2) delivered from a high resolution chromatographic column. Reponses are indicated as percent change of spikes over the spontaneous activity from the total ORNs within the sensilla. Chain length and the position of methyl group defined the excitatory patterns. Significantly higher activity was recorded for *meta*- and *para*-cresol. N=3; * p <0.05

Finally, we evaluated the behavioral responses of the black legged tick to the electrophysiologically active and related phenols. In general phenols did not elicit robust responses. Overall, a large number of ticks were unresponsive in the set-up to tested stimuli. The data were not normally distributed (Shapiro-Wilk test, *p*<0.05). However, Kruskal–Wallis Rank Sum Test (Kruskal and Wallis 1952) revealed a ***χ***^2^= 30.05, df = 10, *p*-value< 0.05 which allowed us to test it for the Dunn test (Dunn 1961). The pairwise comparisons of overall tick activity using the Dunn test revealed were no significant differences in overall tick behavior (ticks making a choice in general vs not making a choice). Paired tests comparing percent activity in treatment with a control arm revealed that only *p*-cresol at −2 dilution induced significant activity wherein ticks preferred the control side (no odor) (Figure 4).

**Figure 4.**
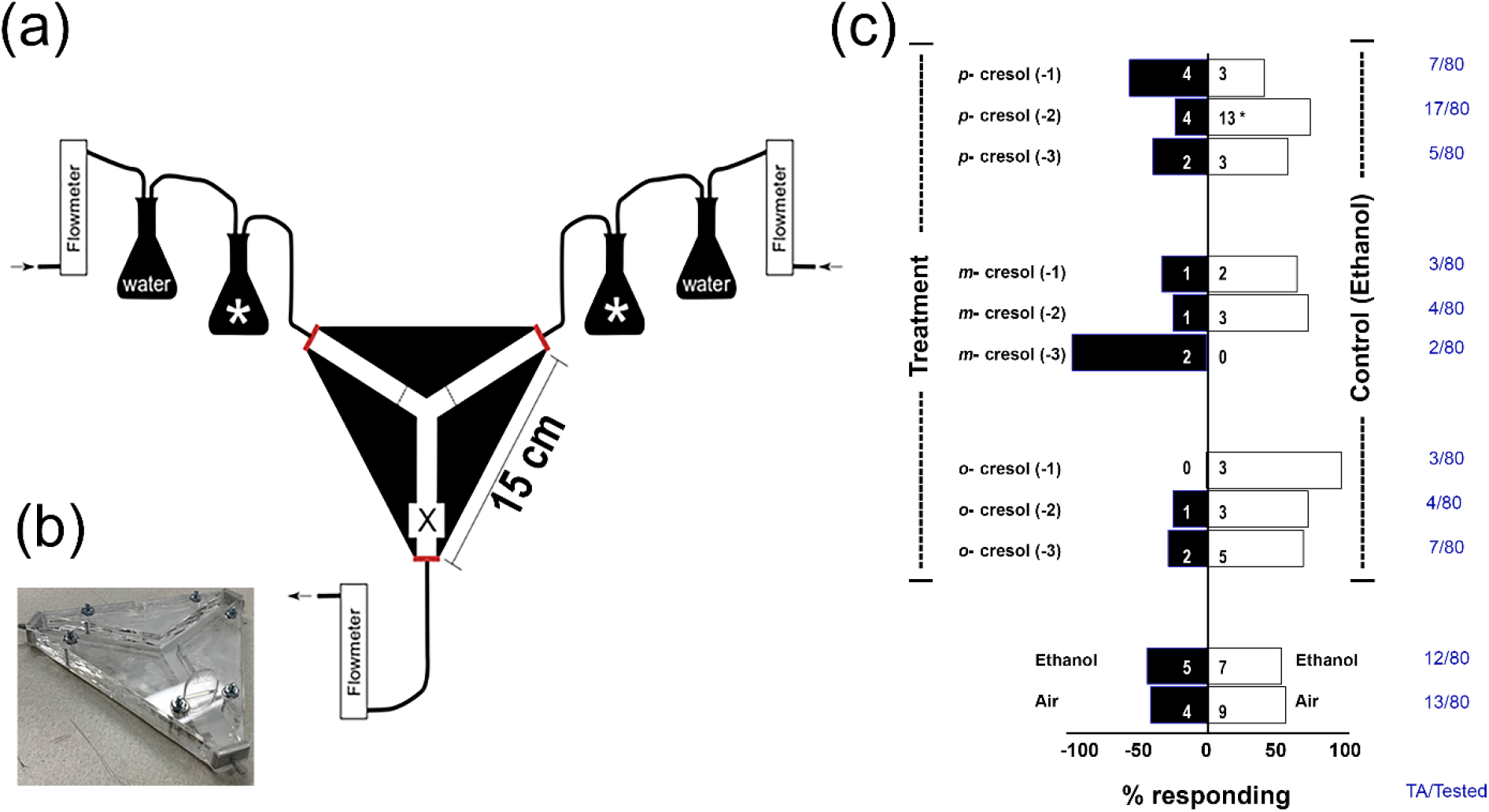
Measuring *I. scapularis* behavioral responses to the electrophysiologically active phenols. (a) Schematic of the dual choice olfactometer. Arrows represent air flow direction. X represents the release point where ticks were placed in the olfactometer at the start of the experiment. Erlenmeyer flasks with * represents flasks that were used as either the control or the experimental; (b) photograph of the apparatus. (c) Behavioral responses induced by methylphenols. Each chemical was tested at three dilutions (1, 10 or 100 μg/μl dilutions represented as −3, −2 or −1 respectively) on 80 adult females. All the chemicals were dissolved in ethanol, that served as control. Right column, in blue, denotes the total number of ticks activated (TA) over the total number tested.

## DISCUSSION

Our electrophysiological experiments present the first SSR and GC-SSR recordings from *I. scapularis*. We found that the large wall pore-single walled (wp-sw) sensillum in the capsule aperture of the Haller’s organ detects a variety of semiochemicals (Figure 1). Phenols, especially the *meta*- and *para*-cresols elicited extreme excitation from an ORN (Figure 2 and 3). Phenols have a special role in the life cycle of many tick species. Since the fascinating discovery of 2,6-dichlorophenol – the first discovery of a chlorinated organic compound in any land animals – as a female sex pheromone in *A. americanum* (Berger 1972), and the later identification of an array of phenols derived from tick extracts (Wood et al. 1975), these chemicals appear in multiple contexts related to tick chemical ecology and sensory physiology.

The neurophysiological response by *I. scapularis* adults to phenols is significant because these compounds are found in the urine (Jemiolo et al. 1995) and forehead secretions of the white-tailed deer (Gassett et al. 1997), one of the main hosts of *I. scapularis*. Additionally, white-tailed deer urine and its derivatives, and semiochemicals derived from the deer’s tarsal or interdigital glands induced varying behavioral responses, including arrestment in this species of tick (Carroll et al. 1996, Carroll 2000). Our chemical analysis of these products revealed phenols as major chemical constituents (Supplemental Figure 1), comparable to the phenolic abundance in the comprehensive analysis of the cattle urine (Bursell et al. 1988). Phenols also act as strong chemostimuli in the obligatory hematophagous tsetse flies, *Glossina* spp. (Hassanali et al. 1986, Bursell et al. 1988) and mosquitoes (Cork and Park 1996).

Interestingly, most of the identified and defined sex pheromones in ticks are substituted phenols, such as *ortho*-nitrophenol and 2,6-dicholorphenol (2,6-DCP) (Sonenshine 2006) that are detected by the olfactory sensilla in ticks, such as in *A. americanus* (Haggart and Davis 1981), *Rhipicephalus appendiculatus* (Waladde 1982), *B. microplus* (De Bruyne and Guerin 1994), *I. ricinus* (Leonovich 2004), *A. variegatum* (Steullet and Guerin 1994b) and *A. cajennese* (Soares and Borges 2012). In most of these studies, ORNs from more than one sensillum responded to the pheromone (s) (Leonovich 1990). This raises many interesting questions in regards to the molecular basis of the detection and perception in ticks (Mulenga 2013). Recent insights into the foreleg transcriptome of the *I. scapularis* offer insights: there were no odorant receptors (ORs) present, however a variety of gustatory and ionotropic receptors (GRs and IRs) were apparent (Josek et al. 2018b). Lack of ORs however does not preclude the ticks from sensing semiochemicals, since phenols, carboxylic acids and aldehydes we found here to be eliciting electrophysiological activity (Table 1) are detected by multiple IRs (Rytz et al. 2013, Gomez-Diaz et al. 2018). Carbon dioxide, a key chemostimulus in tick biology (Wilson et al. 1972, Eisen and Paddock 2020) is detected by GRs in insects (Jones et al. 2007). However, in *I. scapularis*, the GRs responsible for carbon dioxide detection are unknown as none of the insect GRs used in detecting carbon dioxide were related to the *I. scapularis* GRs (Josek et al. 2018b). In addition to the molecular anomalies, neuronal projections from the Haller’s organ do not appear to follow the 1:1 rule of 1 OR/ORN for each antennal glomerulus (Šimo et al. 2014, Borges et al. 2016), generally accepted in a variety of insects (Jefferis 2005, Vosshall and Stocker 2007).

Behavioral responses induced by the biologically active and other phenols were not so clear from our bioassays. Similar results – lack of any significant behavioral activity – to the biologically active phenols was observed for *I. ricinus* (Leonovich 2004). The dynamics of phenols production and reception is complex in ticks. In two species of *Rhipicephalus* (*R. appendiculatus* and *R. pulchellus*) phenols were produced by females only while feeding, and the highest attraction was measured only from adult males (Wood et al. 1975). Our analysis of the commercial odorous deer gland extracts by GC-MS also revealed an abundance of phenol and its substitutes, further reinforcing the view that arthropods, in general, use a limited array of chemicals parsimoniously (Blum 1996, Syed 2015). Finally, we note that behavioral output is also the summation of the inputs from multiple sensory modalities (Barrozo 2019). Therefore, a detailed electrophysiological analyses of all the ORNs from the tick tarsi will provide a comprehensive understanding of the tick’s chemical landscape thus providing us a rich repertoire of chemostimuli. The olfactory sensilla in ticks have been described only from the forelegs, and none elsewhere on the body such as other legs, mouthparts, or on the idiosoma (Hess and Vlimant 1986). This comprehensive morphological analyses of the foreleg tarsal sensory structures and their neuronal output from *A. variegatum*, *A nuttalli*, *D. marginatus*, *B. microplus*, *I. ricinus* and *O. moubata* estimated *ca*. 30% of the tarsal output neurons as ORNs (Hess and Vlimant 1986).

In conclusion, by studying chemorception in the Haller’s organ of *I. scapularis* from multiple viewpoints – the odorants it can detect and how the tick respond to them behaviorally – we have started to assemble a more complete view of how these ticks are able to detect and discriminate hosts and mates. An understanding of the olfactory structures – peripheral and central – and their function (Šimo et al. 2014) will be critical to the development of tools to develop attractants and repellents (Bissinger and Roe 2010, Syed 2014, Carr and Salgado 2019). Our physiological and behavioral analyses of this important disease vector will aid in in designing novel management strategies towards suppressing *I. scapularis* populations to reduce the disease burden.

## ACKNOWLEDGEMENTS

We thank Drs. Brian F. Allan, Allison Parker, Erin Allman-Updyke, Elijah Juma and L. Page Fredericks, members of the Syed laboratory, Drs. Paul Hickner, Omprakash Mittapalli and Ms. Pritika Pandey, for their comments on the early draft of this manuscript. This research work was supported by funding from National Institute of Food and Agriculture, US Department of Agriculture to ZS (under HATCH Project 2353077000) and the University of Illinois at Urbana-Champaign School of Integrative Biology’s Harley J. Van Cleave Research Award and Philip W. Smith Memorial Fund to TJ.

## Notes

### Competing Interest Statement

The authors have declared no competing interest.

