## Supplemental Figure 1 for "Neurophysiological and Behavioral Responses of *Ixodes scapularis* to host odors"

**SUPPLEMENTAL MATERIAL**

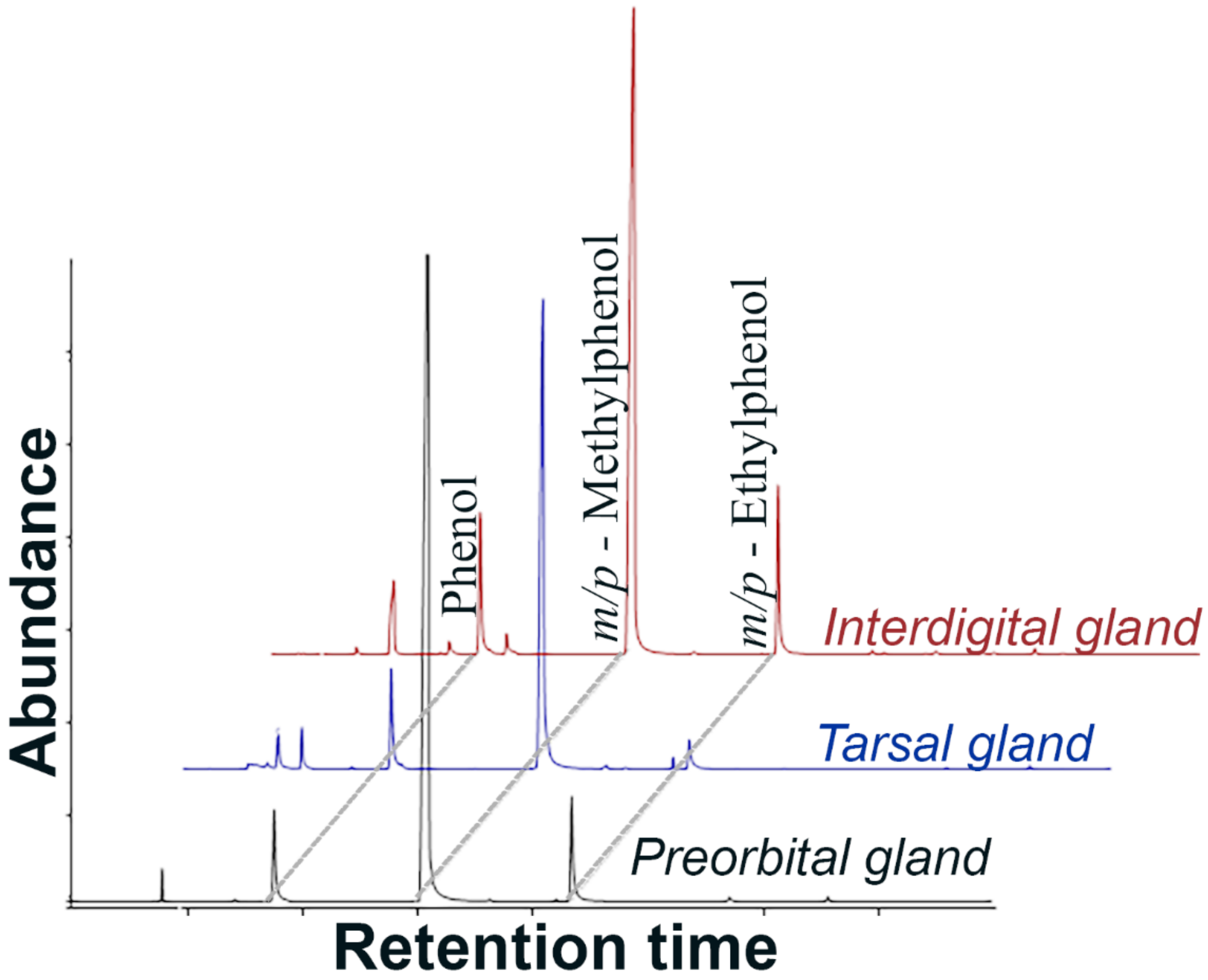


Supplemental Figure 1. GC-MS analyses of the commercial deer gland derivatives reveal abundance of phenols. Total Ion Chromatograms (TIC) are slightly adjusted on time scale for clarity; dotted lines across the three TIC highlight the three major constituent peaks. The calculated Kovat's Indices (KI) are 988, 1080 and 1168 respectively (from left to right) on DB-5 column.
